# Dorsolateral striatum, not motor cortex, is a bottleneck for responding to task-relevant stimuli in a learned whisker detection task in mice

**DOI:** 10.1101/2022.03.03.482906

**Authors:** Behzad Zareian, Angelina Lam, Edward Zagha

## Abstract

A learned sensory-motor behavior engages multiple brain regions, including the neocortex and the basal ganglia. How a target stimulus is selected by these regions remains poorly understood. Here, we performed electrophysiological recordings and pharmacological inactivations of motor cortex and dorsolateral striatum to determine the representations within and functions of each region during performance in a selective whisker detection task in male and female mice. From the recording experiments, peak pre-response activity and significant choice probability emerged in the motor cortex before the dorsolateral striatum, suggesting a sensory-to-motor transformation in which the striatum is downstream of motor cortex. We performed pharmacological inactivation studies to determine the necessity of these brain regions for this task. We found that suppressing the dorsolateral striatum, but not motor cortex, severely disrupts responding to task-relevant stimuli, without disrupting the ability to respond. Together these data support the dorsolateral striatum, and not motor cortex, as an essential node in the sensory-to-motor transformation of this whisker detection task.

**Significance Statement:** We learn to do various sensory-motor behavior in our daily life, such as clicking on a journal article that looks interesting, among other articles. There are parts of our brain that are active when we do these learned behaviors, such as motor cortex and basal ganglia. But what is the order of activation of these regions? Which of them is necessary for responding to task-relevant sensory information? To answer these questions, we trained mice in a whisker-based target selection task and used recording of neural activity and inactivation of subregions within motor cortex and basal ganglia in expert mice. Our findings show dorsolateral striatum, a region within basal ganglia, is a bottleneck for performing task-related sensory-to-motor transformation.

## Introduction

Goal-directed behavior requires the ability to selectively respond to target sensory stimuli (selection), while inhibiting responses to extraneous or distractor stimuli. In simple Go/NoGo tasks, sensory selection involves transforming sensory responses into motor commands. Neuronal recording studies have demonstrated full sensory-to-motor transformations unfolding across the neocortex, with motor planning and motor command signals present most robustly in motor cortices (de Lafuente & Romo, 2006; Esmaeili et al., 2021; Finkelstein et al., 2021; Hanes & Schall, 1996; Inagaki et al., 2019; Li et al., 2015; Li et al., 2016; Moran & Desimone, 1985; Salinas & Romo, 1998; Siegel et al., 2015). These studies provided strong motivation for considering the neocortex as the primary structure implementing goal-directed, sensory-to-motor transformations (Figure 1C). Simultaneously, anatomical, physiological, and lesioning studies of the basal ganglia identified these subcortical structures as essential for implementing action selection and initiation (Figure 1D) (Alexander & Crutcher, 1990; Bergstrom et al., 2020; Bergstrom et al., 2018; Graybiel et al., 1994; Grillner et al., 2005; Jin & Costa, 2010; Stephenson-Jones et al., 2011; Yin et al., 2004). Motor cortex and dorsolateral striatum are heavily interconnected, as determined by both anatomical and functional studies (Foster et al., 2021; Frank et al., 2001; Gordon et al., 2022; Hintiryan et al., 2016; Hunnicutt et al., 2016; Kupferschmidt et al., 2017; Lee et al., 2019; McGeorge & Faull, 1987; Peters et al., 2021; Saunders et al., 2015), and therefore sensory selection may be mediated by the coordinated activities of both regions. Traditional models of cortico-striatal coordination propose that motor plans are generated in motor cortex and then selected in the basal ganglia. The output of this striatal selection is propagated via the thalamus back to motor cortex, which ultimately sends out the motor command (Figure 1E) (Hoover & Strick, 1999; Middleton & Strick, 2000; Redgrave et al., 1999). And yet, the basal ganglia project to multiple regions besides the neocortex which may also trigger motor commands (Figure 1F) (Guo et al., 2017; Mink, 1996; Utter & Basso, 2008; Yin & Knowlton, 2006). Revealing the functional organizations of the motor cortex and the basal ganglia requires conducting representational and causal studies from both structures in subjects performing the same behavioral task (Antzoulatos & Miller, 2011; Brockett et al., 2022; Clarke et al., 2008; Kupferschmidt et al., 2017; Muhammad et al., 2006; Pasupathy & Miller, 2005; Peters et al., 2021; Pimentel-Farfan et al., 2022)

**Figure 1.**
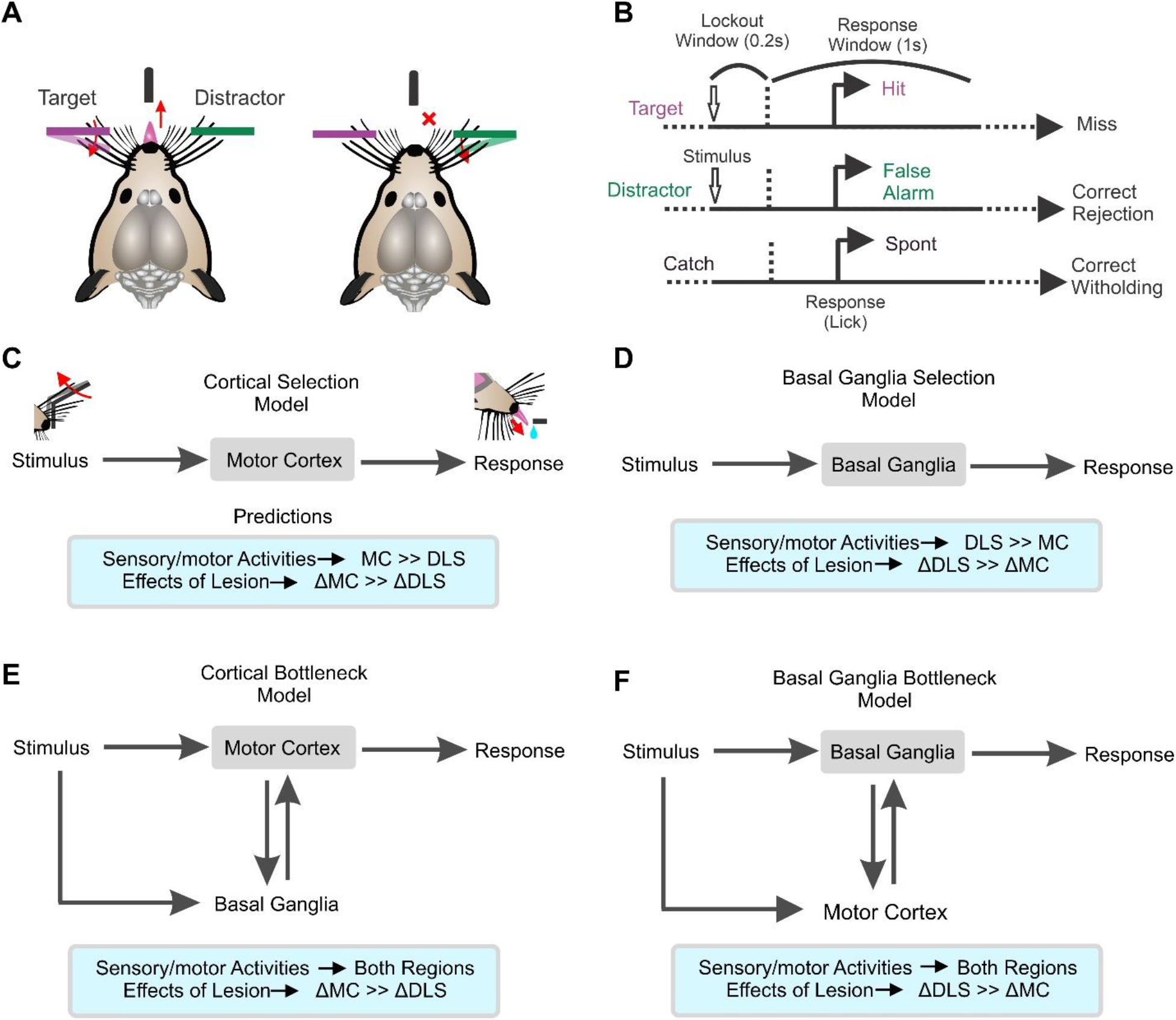
Behavioral task, trial structure and possible functional organizations underlying sensory selection. (A) Illustration of the behavioral task. Mice learn to respond (lick) to small, transient whisker deflections in one of their whisker fields (purple - target) and to withhold licking to identical deflections in their opposite whisker field (green - distractor) (modified from (Aruljothi et al., 2020)). (B) Diagram of task structure for the operant, selective detection task (also see methods). Following a variable (5-9s) intertrial interval, mice receive either a target stimulus, distractor stimulus, or catch trial. After stimulus onset, a lockout window of 0.2s is implemented in which responding is punished by immediately restarting the intertrial. Responses during the response window of target trials are considered hits and responses during the response window of distractor trials are considered false alarms. (C-F) Different models (top) and their experimental predictions (bottom) for the pathways involved in converting whisker stimuli into motor responses. The models differ in the involvement and importance of sensory-to-motor transformations through the motor cortex versus the basal ganglia.

The mouse whisker system is an ideal model system to test this functional organization, due to its simplified and well-characterized neural anatomy. Whisker stimulus responses propagate to the contralateral whisker representation of primary somatosensory cortex (S1). In turn, S1 projects directly and robustly to sub-regions of both the motor cortex (MC) and the dorsolateral striatum (DLS) (Mao et al., 2011). This study focuses on these sub-regions of MC and DLS, as possible signaling pathways for sensory selection. Prior representational and causal studies implicate the involvement of both MC and DLS in whisker detection/discrimination tasks (Aruljothi et al., 2020; Guo et al., 2014; Hong et al., 2018; Lee et al., 2019; Li et al., 2015; Li et al., 2016; Sippy et al., 2015; Zagha et al., 2015; Zareian et al., 2021). And yet, the functional organizations between these regions in the context of whisker sensory selection remains poorly understood. In this study, we directly compared the representational and functional properties of MC and DLS in a Go/NoGo selective whisker detection task (Figure 1). We previously demonstrated robust sensory, motor, and choice signals in whisker-related MC, identifying this region as the most likely site of sensory-motor transformations within dorsal neocortex for this task (Aruljothi et al., 2020; Zareian et al., 2021). In the current study, our data support a functional organization in which DLS is downstream of MC and is an essential bottleneck for transforming whisker stimuli into motor commands.

## Methods

### Animals

Experiments performed in this study were approved by the Institutional Animal Care and Use Committee of University of California, Riverside. Male and female mice of two strains were used for the experiments: wild type (C57BL/6J) and Thy1-ChR2 mice (all purchased from The Jackson Laboratory or bred in our own colony). Data from mice of each sex and strain were combined, and the transgenic feature was not exploited in these studies. All mice were housed on a 12h light/12h dark cycle. Food was always accessible to mice outside of the behavioral training sessions.

### Surgery

All experiments were performed on head-fixed mice. To attach the headpost to the skull, the mice were first anesthetized with a mixture of ketamine (100 mg/kg), xylazine (10 mg/kg), and isoflurane (1-2%) throughout the surgery. They were additionally administered meloxicam (5 mg/kg) and enrofloxacin (5 mg/kg) at the day of the surgery and for two days post-surgery. A 10 mm × 10 mm part of scalp was resected. A stainless steel headpost with length of 3 cm and a weight of 1.5 grams with a central window of 8 mm × 8mm, was attached to the skull using cyanoacrylate glue. The 8 mm × 8 mm exposed window was sealed with Kwik-sil. After the surgery, the mice recovered on a heating pad. Behavioral experiments were started a minimum of 3 days after recovery from surgery. At the day of recording or inactivation, small craniotomies (~0.5 mm) were made under isoflurane anesthesia to access the relevant cortical and subcortical regions (see below).

### Training

We trained mice in a selective detection task, which we have characterized in previous studies (Aruljothi et al., 2020; Marrero et al., 2021; Zareian et al., 2021). The mice were head-fixed in a custom-made setup. Piezo-controlled paddles were placed symmetrically within bilateral whisker fields contacting multiple whiskers. Target and distractor whisker fields were assigned at the beginning of training (target as the right whisker field) and remained constant throughout. Sensory stimuli consisted of small, rapid deflections of either whisker field. The deflections ranged from 0.01-0.2 seconds (s) in duration with a velocity of 10 mm/s, always equal for target and distractor stimuli. For most of the training sessions, two different stimulus amplitudes were used, one (large) near the saturation of the mouse’s psychometric range and the other (small) near the midpoint. Mice reported stimulus detection by licking a central lick port. A lockout of 0.2 s was imposed between stimulus onset and response window, and mice were punished with a timeout for responding during this delay. Following 5-9 s inter-trial intervals, mice received either target, distractor, or catch (no stimulus) trials. All licking outside the post-target response window were punished by time-out (resetting the inter-trial interval). Water rewards for responding during the post-target response window were delivered from the same lick port used to report stimulus detection. Mice were water restricted throughout the training period, with the goal of receiving all water during behavioral trainings. Supplemental water was given in the home cage if weights fell below 85% of initial weights. Behavioral training was implemented using custom MATLAB scripts and Arduino Uno boards to trigger task stimuli and report licking responses. For further details of the behavioral training and training stages, see (Aruljothi et al., 2020).

Mice were considered expert in the task once they achieved a target-distractor discrimination d-prime > 1, for three consecutive days *(d-prime=norminv* (Hit rate)- *norminv* (False alarm rate), in which *norminv* is the inverse of the standard normal cumulative distribution value). All recording and inactivation experiments were performed in animals that had reached expert performance. For the electrophysiological recording sessions, performance values (discrimination d-prime) were as follows: target-aligned MC: 2.7 +/− 0.6, n=11, distractor-aligned MC: 2.5 +/− 0.6, n=19, target-aligned Str: 2.3 +/− 0.6, n=10, distractor-aligned Str: 2.3 +/0.7, n=11 (mean +/− STD). Performance of the mice during muscimol inactivation are reported in the Results, since behavioral performance was the dependent variable under examination.

### Electrophysiological recordings

20 mice were used for electrophysiological recording experiments (see Table 1 for the number of sessions used in each analysis). Recordings were conducted following at least 20 minutes after recovery from isoflurane anesthesia. Recovery was assessed based on normal mouse behavior within their home cage and high engagement during the first few minutes of the task. For MC and DLS recordings, Neuronexus laminar probes with 16 sites and 100 μm spacing were used (A1×16-5mm-100-177-OA16LP or A1×16-5mm-100-177-A16). For each recording, the probe was advanced slowly in the brain using hydraulic Narishige micromanipulators. Electrophysiology data were acquired using Neuralynx recording system and Cheetah viewer software. The data were acquired at 32 kHz, then subsequently filtered at 600 to 6000 Hz for spike analyses.

**Table 1.**
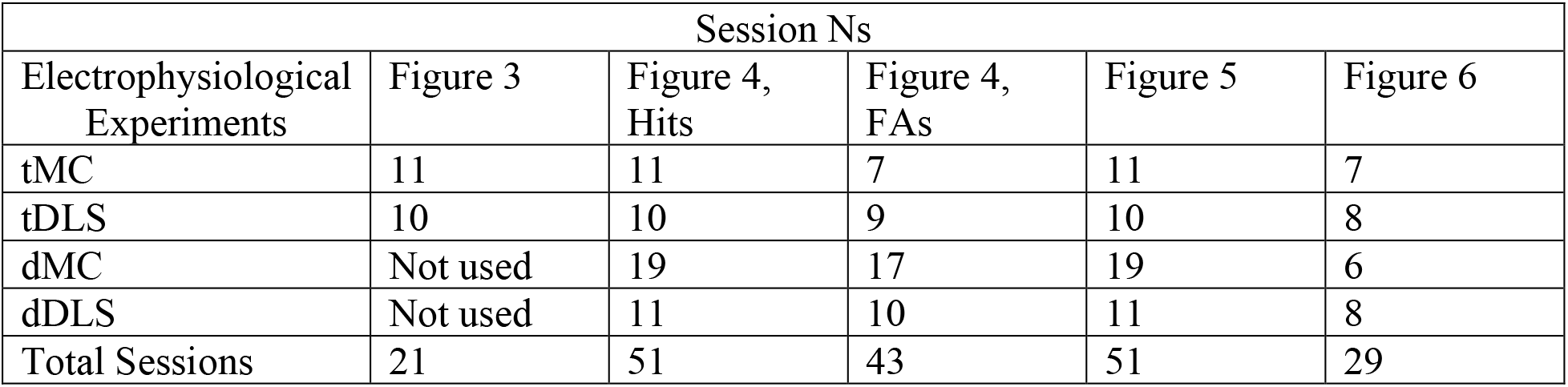
Session number (N) used for electrophysiological recordings. The number of sessions for each region used for analyses in different figures, related to recording experiments.

| Session Ns |  |  |  |  |  |
| --- | --- | --- | --- | --- | --- |
| Electrophysiological Experiments | Figure 3 | Figure 4, Hits | Figure 4, FAs | Figure 5 | Figure 6 |
| tMC | 11 | 11 | 7 | 11 | 7 |
| tDLS | 10 | 10 | 9 | 10 | 8 |
| dMC | Not used | 19 | 17 | 19 | 6 |
| dDLS | Not used | 11 | 10 | 11 | 8 |
| Total Sessions | 21 | 51 | 43 | 51 | 29 |

Craniotomies for MC recordings were centered on 1 +/− 0.5 mm lateral, 1 +/− 0.5 mm anterior to bregma. These MC coordinates were chosen to target the S1 projection zone, as in our previous studies (Zareian et al., 2021). The probe was advanced until the last site was barely visible at the surface (16 sites spanning 1.5 mm within cortex and below). At these coordinates, the thickness of layer 1, 2/3 and 5A of MC collectively span the superficial (dorsal) 500-600 μm (Hooks et al., 2013) (also see Figure 2C). Therefore, we considered recording sites within this range as ‘superficial MC’ and the more ventral recording sites as ‘deep MC.’ DLS coordinates were as follows [from bregma]: 2.5 +/−0.5 mm lateral, 0.7 +/− 0.4 mm posterior (also see Figure 2F). The probe was advanced deep inside the DLS (distal tip 2300-2500 μm below the pial surface). The coordinates were initially chosen based on Allen Brain Institute Mouse Connectivity atlas, selecting the region within DLS receiving the highest density inputs from S1.

**Figure 2.**
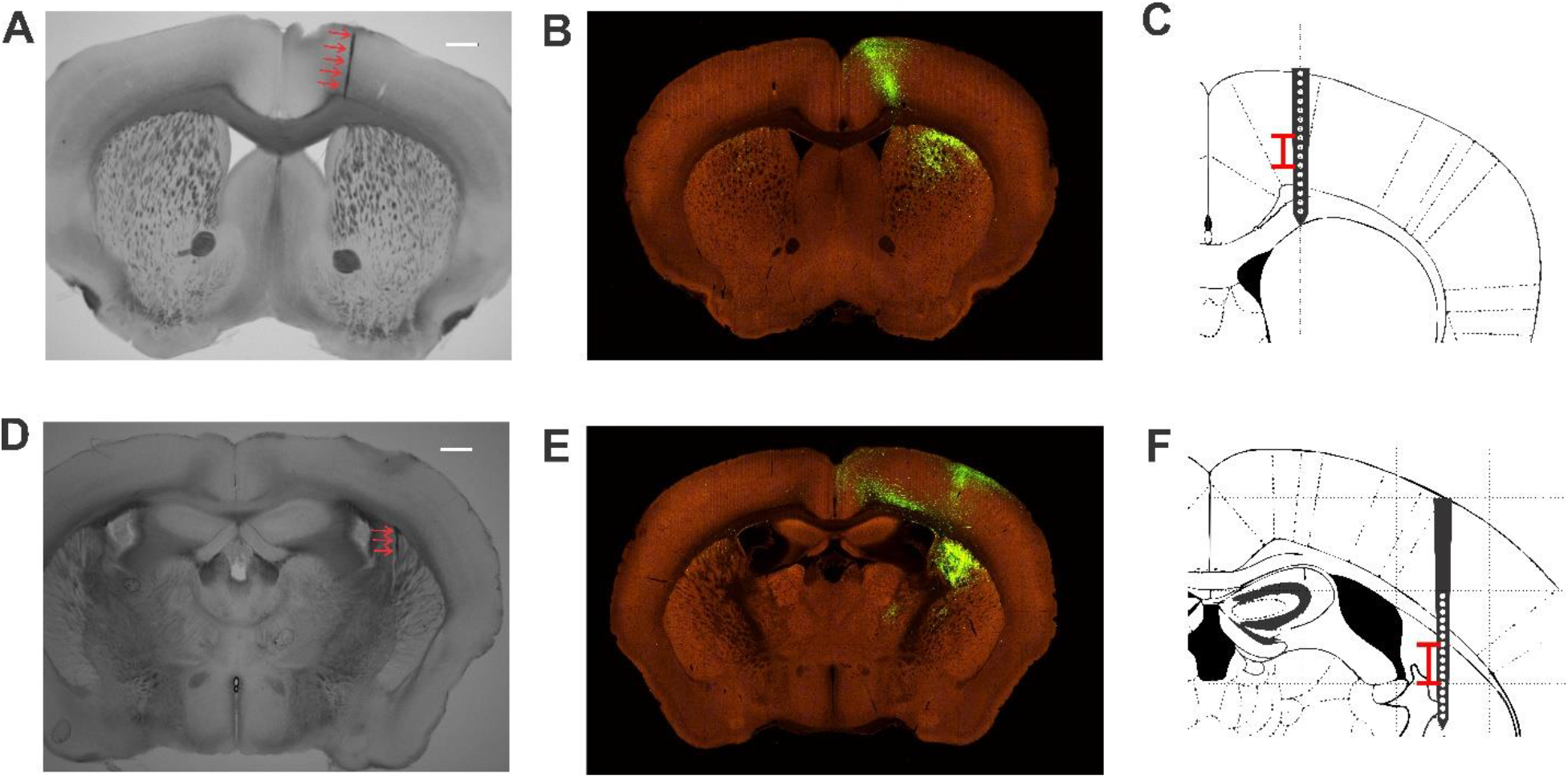
MC and DLS sub-region targeting. (A, D) Coronal brain sections depicting the needle tracts (red arrows) from targeting MC (A) and DLS (D). (B, E) Findings from whisker-related S1 anterograde labeling from the Allen Brain Atlas: Mouse Connectivity (experiment 126907302), showing the S1-projection sites (green) in MC (B, section 49) and DLS (E, section 62). (C, F) Schematics of the same coronal sections as in [A, B] and [D, E], respectively, illustrating the coverage of the laminar multi-site electrode recordings. Red bars indicate the mean +/− 1 standard deviation for the sites with maximum sensory encoding within each region. Scale bars in [A] and [D] are 1 mm.

### Muscimol Inactivation

Injections of 2 mM muscimol in normal saline were performed using the Nanoject III Programmable Nanoliter Injector from Drummond Scientific Company fitted with a borosilicate glass micropipette. The same surface coordinates were used for muscimol injections as described above for electrophysiological recordings. For MC injections, 250 nL of muscimol was injected at a rate of 1-3 nL/s at 1 mm deep (from the pial surface) and another 250 nL was administered at 0.5 mm deep, for a total of 500 nL, to target both superficial and deep layers of MC. Based on previous experience using nearly identical protocols, this application causes inactivation of cortex approximately 1 mm in diameter as observed from the dorsal surface [see Figure 4D in (Salkoff et al., 2020)]. For DLS injections, a single bolus of 250 nL of 2 mM muscimol was injected at a rate of 1-3 nL/s at 1.7 mm deep. Based on the descriptions above, we expect this protocol to inactivate a volume of the striatum less than 1 mm in diameter.

For muscimol inactivation studies, expertly performing mice were assigned to alternating control performance days, without muscimol exposure, and experimental performance days, with muscimol inactivation. Sites for muscimol inactivation were randomized for each mouse, such that the order of inactivation varied (both within and across mice). For daily behavioral testing (also see Figure 7A), mice were first placed in a classical conditioning version of our task (see (Aruljothi et al., 2020)) for 2 minutes to ensure licking responses were intact when presented with water reward cued by the opening of a solenoid. Then mice were tested in the full selective detection task for 1 hour. After 1 hour of testing if mice collected fewer than 10 rewards, they were considered non-performers. 10 rewards is 2-3 standard deviations below the number of rewards achieved by non-injected control mice within the same duration (62 +/− 21 rewards [mean +/− SD]). Non-performing mice were placed back in the classical conditioning task for 15 minutes to again assess for licking to solenoid-cued water rewards. If mice were still performing the full selective detection task after 1 hour, they were permitted to perform that task until unmotivated (as determined by time since previous reward greater than 10 minutes). The threshold for consideration as ‘performing’ in the classical conditioning task was responding to >50% of rewards, in either testing phase.

### Histology

Verifications of silicone probe and muscimol injection sites were performed on wildtype C57BL/6J mice not previously utilized for muscimol inactivation experiments. A borosilicate glass micropipette was inserted into MC or DLS using similar protocols as for muscimol inactivation studies. One hour after injection, mice brains were processed to visualize the pipette tract. Mice were sacrificed using Euthasol euthanasia solution and transcardially perfused with 20-40 mL of 1X Phosphate-Buffered Saline solution (PBS) followed by 20-40 mL of 4% Paraformaldehyde (PFA) in 1X PBS before the brain was dissected and stored in 4% PFA overnight. The following day, brains were rinsed with 1X PBS three times, then embedded in a 3% agarose in 1X PBS solution. 120 μM slices were collected using a Leica VT1000 S vibrating blade microtome and mounted onto glass slides using glycerol and a coverslip. 2X images were collected using a Keyence BZ-X710 microscope under brightfield illumination (Figure 2A and D).

Comparison tracing studies (Figure 2B and E) are from the Allen Brain Mouse Connectivity Atlas, experiment 126907302. For the schematics depicting approximate recording sites (Figure 2C and F), we traced the outlines of slices from The Mouse Brain in Stereotaxic Coordinates (−1.06 and 0.98 mm from Bregma for DLS and MC, respectively) (Paxinos & Franklin, 2019). We compared the cortical thickness from the atlas to our functional estimates based on white matter location and determined a scale factor of 17%. Accordingly, a 17% reduction was applied to the mapping of the recording sites onto the atlas schematics.

### Quantification and statistical analysis

Analyses were performed using custom MATLAB scripts or SPSS and displayed using Corel Draw. For all statistical analyses, we used alpha=0.05 as significant threshold, unless otherwise stated. Data are presented as mean +/− standard error of the mean, unless otherwise stated. Analyses focused on large amplitude target and distractor stimuli.

### Multiunit activity (MUA) analyses

Behavioral and combined behavioral-recording sessions were truncated to include a single engaged period (a continuous bout of at least 10 minutes of task performance with no gaps in responding (licking) greater than 60 seconds). MUA was identified as negative-going threshold crossings over 3×standard deviation of the band-pass filtered voltage fluctuations throughout that session.

For MC recordings we used linear multielectrode arrays with 100 μm spacing. Channels 1 to 7 (spanning the superficial 600 μm) were considered as superficial layers. The remaining channels (8 to 16) were considered as deep layers. For DLS data analyses, we first determined the location of the white matter, as the site within the middle 5-12 sites with the minimum average spiking. The recording sites below that were considered as putative DLS sites.

Spike counts were binned in 5 ms bins throughout each recording session and combined as needed for larger window analyses. We calculated sensory detection and choice probability values using signal detection theory (Zareian et al., 2021). Briefly, the sensory detection was calculated by considering the area under receiver operating characteristic (ROC) curve constructed from plotting cumulative distributions of spiking of a post-stimulus window against pre-stimulus baseline activity. For baseline activity, three consecutive epochs with the same size as the post-stimulus window were considered. Post-stimulus window sizes were either 5 ms bins for continuous d-prime traces (Figure 4 A-D) or a single 100 ms window immediately post-stimulus (Figure 4 E and F). For latency to reaction time analyses, a window of 1 s before the reaction time was considered as baseline.

Similarly, choice probability was calculated as the area under the ROC curve constructed from plotting the cumulative distributions of spiking activity on hit trials against miss trials. The choice probability traces were calculated from 50 ms sliding windows with 90% overlap. For statistical comparisons, the session choice probability values were compared to chance level (50%). For choice probability analysis, sufficient hit and miss trials were assessed based on previously described criteria (Zareian et al., 2021). Choice probability latency was determined as the time from stimulus onset to rise above 60% choice probability.

### Muscimol inactivation analyses

For the muscimol inactivation experiments, we obtained task engagement time, hit rate, false alarm rate, d-prime, and criterion using a 1-hour time window of selective detection task performance (Figures 7, 8). Data were averaged across all sessions and all mice for each set of analyses (from n=7 total mice, see Table 2 for the number of sessions used). Task engagement times (Figure 7D) were calculated as the sum of all times mice were active during the task, as defined by no lapse in licking greater than 1 minute.

**Table 2.** Session number (N) used for muscimol inactivation. The number of sessions for each region used for analyses in different figures, related to control and inactivation experiments.

| Session Ns |  |
| --- | --- |
| Inactivation Experiment Type | Figures 7, 8 |
| Control | 72 |
| tMC | 12 |
| tDLS | 10 |
| dMC | 13 |
| dDLS | 10 |
| Total Sessions | 117 |

### Statistical analyses

For statistical comparisons of recording experiments, t-test or ANOVA were used. ANOVA statistics was calculated using either SPSS or MATLAB. We considered region and hemisphere effects by running 2-way ANOVA tests; performance measures were grouped regionally (main effect of MC vs. DLS regions regardless of the hemisphere) and grouped by hemisphere (main effect of target-aligned vs. distractor-aligned regions regardless of brain regions). ‘Target-aligned’ or ‘distractor-aligned’ refers to the hemisphere contralateral to the whisker field that receives target or distractor paddle deflections, respectively. Interaction effects were also considered (regional × hemisphere). For pairwise comparisons, Tukey-Kramer post-hoc multiple comparison test using *multcompare* function in MATLAB were conducted between pair of conditions (tMC, tDLS, dMC, dDLS). For muscimol inactivation studies, we additionally compared overall performance rates using Chi-square tests and compared inactivation sessions to non-injected control sessions using unpaired t-tests.

## Results

### Behavioral task, model predictions, and regions of interest

We trained mice in a head-fixed, whisker-based selective detection task (Aruljothi et al., 2020; Marrero et al., 2021; Zareian et al., 2021). In this task, mice learned to respond (lick) to small, transient whisker deflections within one whisker field (target) and to ignore identical whisker deflections in the opposite whisker field (distractor) (Figure 1A). Due to the lateralization of the somatosensory system, this task configuration establishes “target-aligned” and “distractor-aligned” cortical and striatal fields that are symmetric across midline and contralateral to the deflected whiskers. In the task structure, we impose a 200 ms lockout between stimulus onset and response window, and mice learn to withhold responding across this delay (Figure 1B). Mice are considered expert in this task once they achieve a separation (d-prime) between hit rate (response to target) and false alarm rate (response to distractor), greater than 1, for three consecutive days (for training details, see Methods and (Aruljothi et al., 2020).

Electrophysiological recording and muscimol inactivation studies were conducted in expert mice, while they were performing the selective detection task. These physiological studies focused on two major outputs of primary somatosensory cortex (S1): the S1-projection subregions of both the motor cortex (MC) and the dorsolateral striatum (DLS) (Mao et al., 2011). Figure 2 depicts examples of MC and DLS targeting, relative to S1-projections.

We recognize four possible functional organizations of these S1-output pathways. If sensory selection is primarily mediated through motor cortex (Figure 1C), we would expect greater sensory and motor activities in MC than DLS, and for inactivations of MC to cause larger impairments of task performance. If selection is primarily mediated through the basal ganglia (Figure 1D), we would expect DLS to show greater sensory and motor activities and larger performance impairments from inactivations. From the cortical bottleneck model (Figure 1E), we would expect robust sensory and motor activities in both MC and DLS, yet inactivating MC to have the larger effects on task performance. From the basal ganglia bottleneck model (Figure 1F), we would expect robust sensory and motor activities in both regions, yet inactivating DLS to have the larger effects on task performance. Subsequent experiments and analyses were designed to distinguish between these models.

### Robust sensory-related and motor-related spiking activities in MC and DLS

We performed laminar electrophysiological recordings and compared the multiunit activity (MUA) signals from MC and DLS. For each recording session, we identified the site with the largest sensory encoding (neurometric d-prime, see Methods for details) and used the activity at that site for all subsequent analyses. For MC, the laminar electrode spanned all layers, and sensory encoding was invariable largest in deep layers (1023 +/− 37 μm, n=30 sessions, n=11 sessions in target-aligned MC and n=19 sessions in distractor-aligned MC). For DLS, sensory encoding was largest 2136 +/− 55 μm from the cortical surface and 452 +/− 53 μm below the putative white matter (n=21 sessions, n=10 sessions in target-aligned DLS and n=11 sessions in distractor-aligned DLS). The approximate locations of these recording sites are shown in Figures 2C and 2F.

In Figure 3, we display post-stimulus and pre-response activities for target stimuli in target-aligned MC (tMC) and in target-aligned DLS (tDLS). In both regions, MUA appeared at short latency after stimulus onset (Figure 3A, B left columns). We consider this activity to be ‘sensory’ because it occurs regardless of trial outcome (Figure 3C, F) and is time-locked to stimulus onset (within the first 100 ms) regardless of the reaction time (Figure 3D, G). We do note differences in activity levels on hit vs miss trials (both pre-stimulus and post-stimulus) (Figure 3C, F), which we analyze further below. To view pre-response activity, we aligned the spiking on each trial to the reaction time (Figure 3A, B right columns). Both tMC and tDLS displayed elevated activity before the response. However, pre-response ramping activity was more evident in tDLS than tMC. This was also appreciated by clustering trials according to reaction time (Figure 3E, H), demonstrating more pronounced transient activations in tDLS within 100 ms before the response (Figure 3H), potentially reflecting motor response triggering. These qualitative observations are followed up with more quantitative analyses below. However, they suggest that both MC and DLS contain robust sensory-related and motor-related activity (consistent with models in Figure 1E, F), with DLS more robustly signaling response triggering.

**Figure 3.**
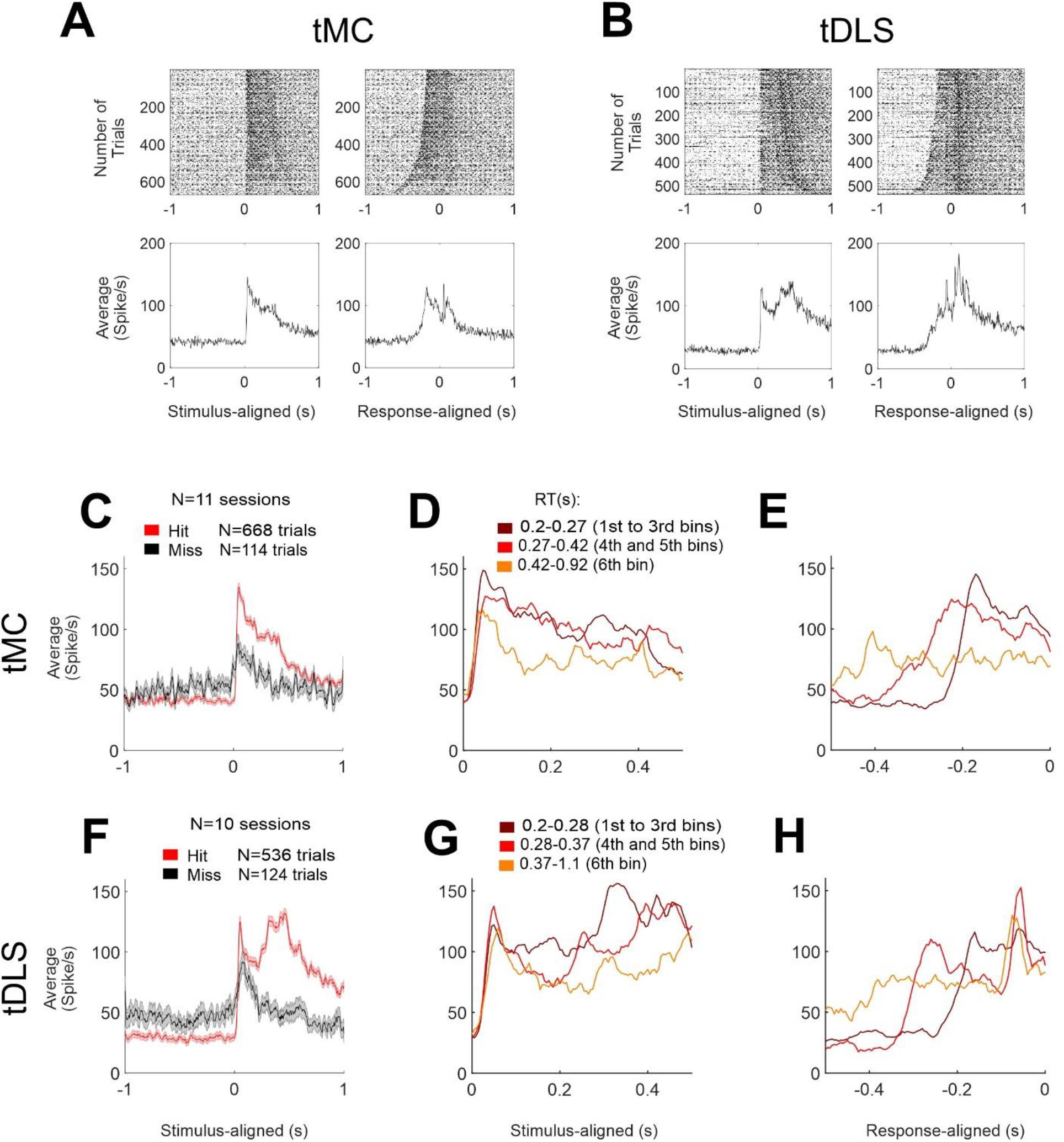
Robust post-stimulus and pre-response spiking activities in target-aligned MC and DLS. (A) MUA recorded from deep layers of target-aligned MC for all hit trials, pooled across sessions. The left column depicts MUA aligned to the target stimulus onset and the right column depicts MUA aligned to the reaction time. Top panels show raster plots sorted based on reaction times from fast (top) to slow (bottom) hits trials. Bottom panels show average of all trials shown in the top panels. Note the rapid post-stimulus (left, putative sensory) and the robust pre-response (right, putative motor) spiking activities. (B) Same as [A] but recorded from target- aligned DLS. (C) Average of all hit (red) and miss trials (black), recorded from deep layers of target-aligned MC (n=11 sessions). (D) Hit trials, pooled across sessions, from target-aligned MC, grouped based on reaction time, aligned to the stimulus onset. Darker colors depict faster reaction time trials. Note the early ‘sensory’ peak, invariant to reaction time. (E) Hit trials, pooled across sessions, from target-aligned MC, grouped based on reaction time (as in [D]), aligned to the reaction time. (F,G and H) Same as [C, D and E] but recorded from target-aligned DLS, also displaying robust sensory and motor alignments.

**Figure 4.**
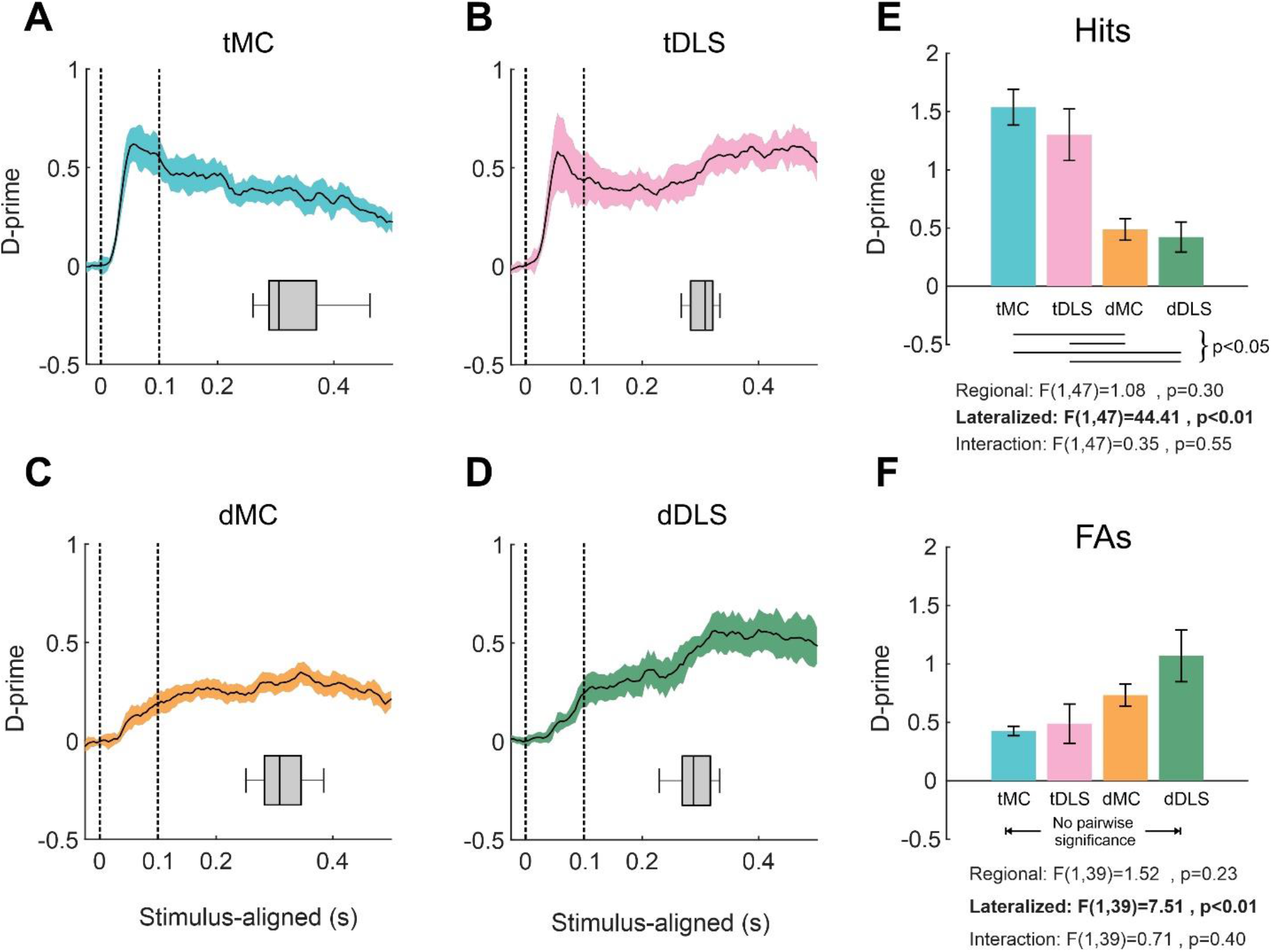
Lateralized sensory responses in MC and DLS. (A) Average peak sensory encoding of target stimulus responses (hits) in deep layers of target-aligned MC. The x-axis denotes time from stimulus onset in seconds. The y-axis denotes neurometric d-prime (as the separation between distributions of pre-stimulus and post-stimulus MUA). Dashed lines indicate the 100 ms post-stimulus window used for quantification in [E]. (B-D) Same as [A], but for target-aligned DLS (B), distractor-aligned MC (C), and distractor-aligned DLS (D). Note the rapid and large increase in d-prime after stimulus onset in target-aligned regions compared to distractor-aligned regions. (E) Sensory encoding of hit trials within each region calculated for the first 100 ms post-stimulus across recording sessions. Lines under the bar graph indicate significant pairwise differences. (F) Same as E, but for distractor stimulus responses (false alarms).

### Lateralized sensory encoding in MC and DLS

In the subsequent analyses, we present data from four brain regions: target-aligned MC (tMC), target-aligned DLS (tDLS), distractor-aligned MC (dMC), and distractor-aligned DLS (dDLS). As such, our main quantification approach follows a 2×2 ANOVA design, assessing for main effects of region (MC vs DLS) or hemisphere (target-aligned vs distractor-aligned).

To quantify sensory encoding in MC and DLS, we computed the neurometric d-prime for each recording session (Figure 4). We limited this analysis only to ‘response’ trials (hits for target trials, false alarms for distractor trials), which presumably contain sensory-motor transformations. On hit trials, we observed that target-aligned regions (both tMC and tDLS) showed large, rapid-onset peaks in their d-prime profiles (Figure 4A, B), compared to more slowly rising sensory encoding in distractor-aligned regions (Figure 4C, D). Averaging across all recording sessions, sensory encoding for hit trials within the first 100 ms post-stimulus (which is 100 ms before the earliest reaction times) peaked at (d-prime): tMC, 1.54 +/− 0.15; tDLS, 1.30 +/− 0.22; dMC, 0.49 +/− 0.09; dDLS, 0.42 +/− 0.13 (Figure 4E). From ANOVA testing, we observed a significant main effect of hemisphere (target-aligned vs. distractor-aligned, F(1,47)=44.41, p<0.01), no significant main effect of region (MC vs. DLS, F(1,47)=1.08, p=0.30), and no significant interaction (F(1,47)=0.35, p=0.55). Such findings indicate sensory encoding on hit trials that is lateralized and similar in tMC and tDLS.

We conducted similar sensory encoding analyses on false alarm trials (Figure 4F). Again, the only statistically significant effect was of hemisphere (target-aligned vs. distractor-aligned, F(1,39)=7.51, p<0.01), yet with larger encoding in distractor-aligned regions. Together, these analyses establish robust and lateralized sensory encoding in MC and DLS, and argue against purely cortical or purely basal ganglia selection models.

### Pre-response peak firing and significant choice probability in MC precedes DLS

Next, we sought to assess the order of activation leading to response triggering. To accomplish this, we analyzed the time course of spiking activity in each region preceding reaction times on hit trials (Figure 5). In all regions, we observed ramping activity preceding reaction times (Figure 5 A-D). However, the time course of these ramping activations differed between regions. We quantified the delay between the 80% percentile of normalized peak neuronal activation and the reaction time (for tMC and tDLS example sessions see Figure 5E). The activation-RT delay for each region was (sec): tMC, 0.18 +/− 0.01; tDLS, 0.11 +/− 0.02; dMC, 0.13 +/− 0.02; dDLS, 0.09 +/− 0.02. Unlike analyses of sensory encoding, activation-RT delay ANOVA testing revealed a significant main effect of region (MC vs. DLS, F(1,47)=9.84, p<0.01) (Figure 5F), with longer delays for MC compared to DLS. We did not observe significant effects of hemisphere or interaction. These analyses suggest that, with regards to response triggering, MC is upstream of DLS.

**Figure 5.**
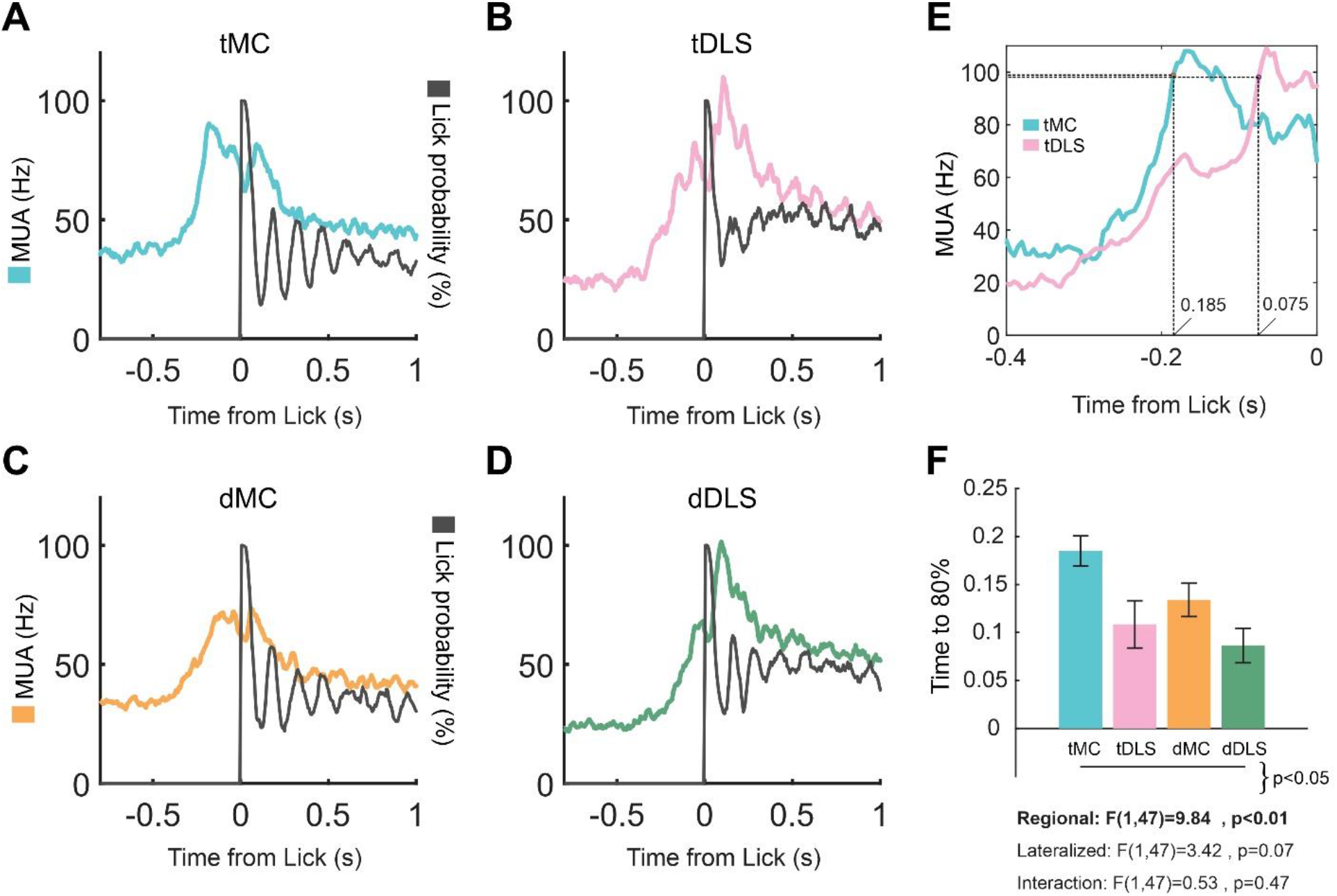
Pre-response peak firing in MC precedes pre-response peak firing in DLS. (A) Average peak multiunit activity aligned to the lick responses on hit trials in deep layers of target-aligned MC. The blue trace shows spike rate (Hz) whereas the black trace shows the lick probability (%). Note the increase in spiking activity before the licking response. (B-D) Same as [A], but for target-aligned DLS (B), distractor-aligned MC (C), and distractor-aligned DLS (D). (E) Average peak multiunit activity for lick responses for example sessions from tMC and tDLS. Dashed lines show the 80% percentile of peak neuronal activation. (F) Delay between the 80% percentile of peak neuronal activation and reaction time across sessions for each recording site. Lines under the bar graph indicate significant pairwise differences.

Next, we used choice probability analyses to gain insights into the temporal latencies of the sensory-motor transformations within each region (Figure 6). Choice probability is the quantification of a relationship between neuronal activity and behavioral outcome, independent of stimulus amplitude (Britten et al., 1996). Above chance (0.5) choice probability indicates epochs in which neural activity is not solely accounted for by sensory processing and may instead reflect decision making and/or motor response processing. In Figure 6A-D, we plot the choice probability time course for target stimuli, as the separation of spiking activity on hit versus miss trials. Target-aligned MC shows prominent early choice probability peaks within 100 ms post-stimulus (Figure 6A). This finding in MC is consistent with our previous report of early choice probability in layer 5 of MC (Zareian et al., 2021). Distractor-aligned MC shows moderate latency choice probability (Figure 6C), whereas target-aligned and distractor-aligned DLS show more gradual rises in choice probability, peaking during the response window >200 ms post-stimulus onset (Figure 6B, D). A comparison of latencies to reach choice probability of 0.6 revealed significantly earlier increases in MC than DLS (sec): tMC, 0.06 +/− 0.02; tDLS, 0.15 +/− 0.04; dMC, 0.11 +/− 0.01; dDLS, 0.20 +/− 0.04; significant main effect of region (MC vs. DLS), F(1,25)=6.97, p=0.01; no significant effect of hemisphere, F(1,25)=2.52, p=0.12, or interaction, F(1,25)=0, p=1.00. These analyses suggest that sensory-motor transformation signaling occurs earlier in MC compared to DLS.

**Figure 6.**
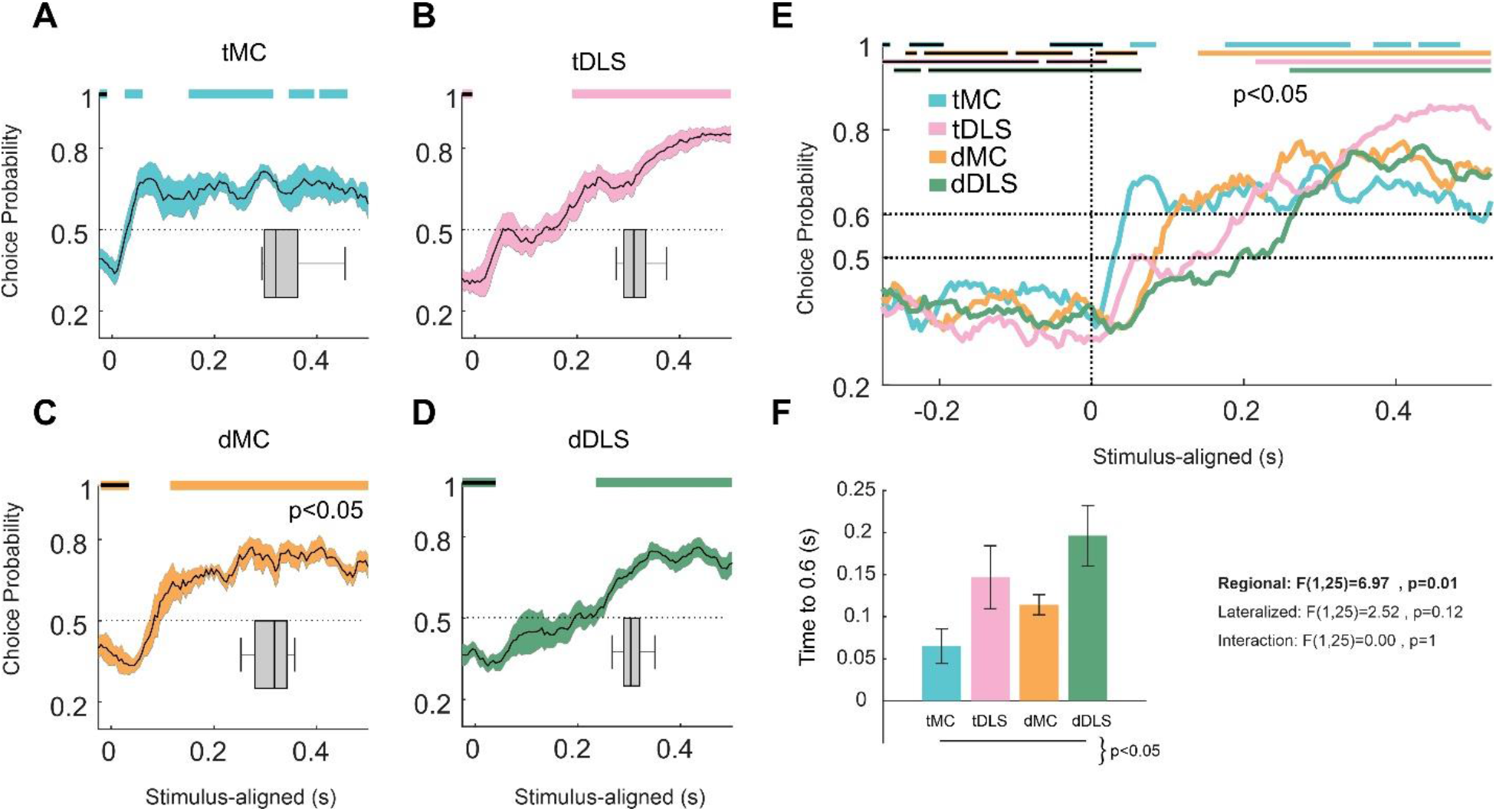
Choice probability in MC precedes DLS. (A) Average choice probability on target trials in deep layers of target-aligned MC. The x-axis denotes time from stimulus onset in seconds. The y-axis denotes choice probability calculated as the separation of spiking activity on hit and miss trials in 50 ms sliding windows with 90% overlap. Significant positive and negative choice probability time points are depicted by plain and filled color bars, respectively, at the top of the panel. Chance level (equal spiking on hit and miss trials) is depicted by the dashed line at 0.5. The grey box-and-whisker plot indicates the reaction time distributions for the same recording sessions. Note the early above chance choice probability, well-preceding the reaction time. (B-D) Same as [A] but for target-aligned DLS (B), distractor-aligned MC (C), and distractor-aligned DLS (D). (E) Choice probability traces, overlapped for all four regions, demonstrating differences in temporal profiles. In addition to chance (0.5), an arbitrary threshold at 0.6 is depicted by a dashed line. (F) Time to reach choice probability of 0.6 across sessions for each recording site. Lines under the bar graph indicate significant pairwise differences.

**Figure 7.**
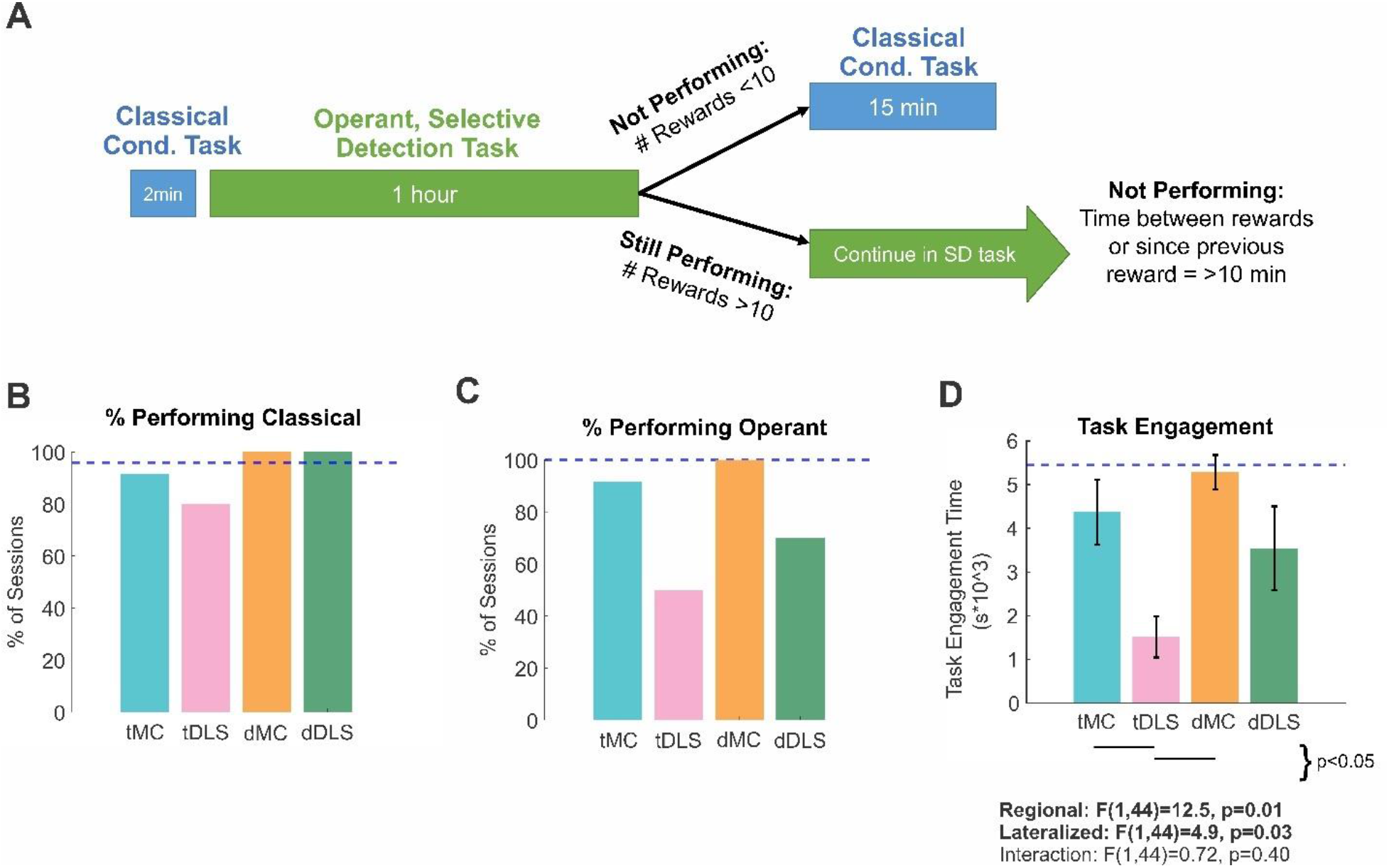
Muscimol inactivation impacts on task performance. (A) Behavioral training workflow for muscimol inactivation experiments (see also Methods). For each behavioral training session, mice were first presented with a classical conditioning task for 2 minutes, followed by the operant, selective detection task for 1 hour. For mice that did not perform the selective detection task, they were again presented with the classical conditioning task for 15 minutes. Mice that did perform the selective detection task continued in this task until unmotivated. (B) Percentage of sessions meeting threshold performance for the classical conditioning task, according to region of inactivation. (C) Percentage of sessions meeting threshold performance for the operant, selective detection task. (D) Length of task engagement within the selective detection task, averaged across all sessions according to the region of inactivation. Lines under the bar graph indicate significant pairwise differences. For [B-D], the horizontal dashed line reflects performance measures from non-injected control sessions.

**Figure 8.**
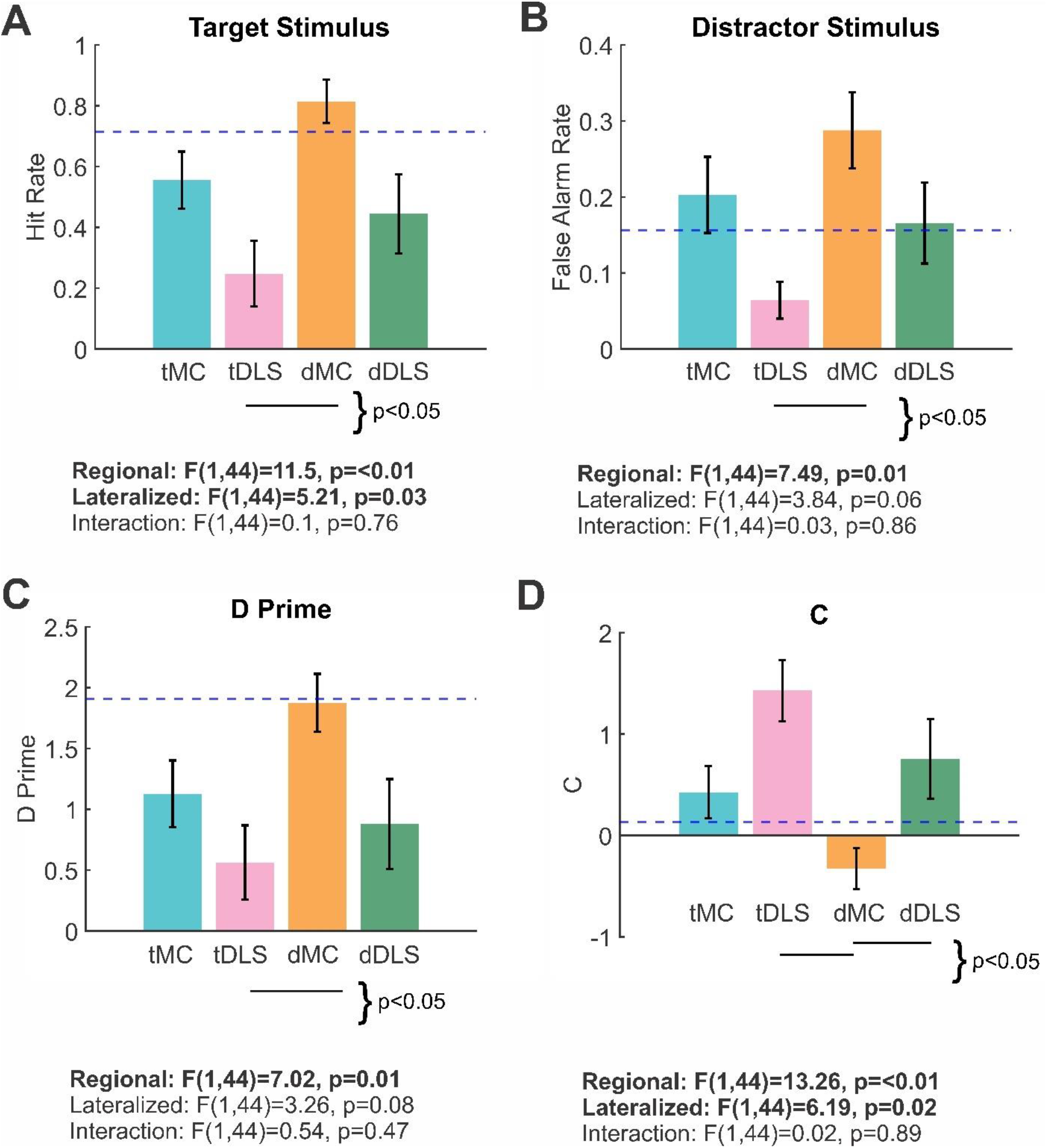
Target-aligned DLS inactivation reduces responding to task-related stimuli. (A) Hit rates within the selective detection task, averaged across all sessions according to the region of inactivation. Lines under the bar graph indicate significant pairwise differences. The horizontal dashed line reflects performance measures from non-injected control sessions. (B-D) Same as [A], but for false alarm rates (B), behavioral d-prime (C), and criterion (D). Note that target-aligned DLS inactivations (pink) caused large decreases in hit rates and false alarm rates, and corresponding increases in the criterion.

We recently reported lower than chance (negative) choice probability before stimulus onset throughout dorsal neocortex, indicating that lower pre-stimulus activity is more likely to result in response (versus no response) outcomes (Marrero et al., 2021). Interestingly, here we observed below chance pre-stimulus choice probability in both MC and DLS (Figure 6A-E). These pre-stimulus baseline differences can also be appreciated in plots of the average spike rates preceding hit and miss trials (Figure 3C and 3F). Thus, noise suppression preceding whisker stimulus detection may generalize to cortical and subcortical structures.

In summary, we find robust sensory, motor, and choice-related signals in both MC and DLS. Sensory signals are lateralized, whereas motor and choice signals are more regionally organized. Notably, we find a temporal ordering of motor and choice signals between regions, suggesting a functional organization with MC upstream to DLS, and DLS activations more directly linked to response triggering.

### Task performance requires target-aligned DLS

Next, we performed muscimol (GABAA receptor agonist) inactivation studies to determine the essential functional contributions of MC and DLS to task performance. Muscimol provides stable inactivation of the exposed region during behavioral testing, enabling assessment of its essential functions that cannot be compensated for on the order of minutes to hours (with full recovery of neural activity within 6 hours, (Arikan et al., 2002)). In mice that had achieved expert performance, we alternated muscimol injected and control (non-injected) testing sessions, randomizing muscimol inactivation among the four sites (tMC, tDLS, dMC, dDLS).

For each behavioral testing session (Figure 7A), we first tested mice in a classical conditioning version of the task in which an auditory cue the mice had previously associated with reward (opening of a solenoid) was presented along with a fluid reward. This was performed to assess global deficiencies in motivation or response initiation/execution. In all inactivation conditions, mice performed this classical conditioning task at a high rate (Figure 7B), with no significant differences between conditions (Pearson Chi-square value = 4.554, p-value = 0.208). These findings argue against global deficits in motivation or response execution. In contrast, we did observe substantial group variance during subsequent performance in the operant, selective detection task (Figure 7C) (Chi-square value = 10.521, p-value = 0.015). Deficits in performance were particularly notable for target-aligned DLS inactivations, in which 50% of the behavioral sessions failed to attain a minimum number of rewards (<10 rewards within 1 hour, compared to an average of 62 rewards for control mice, see also Methods for descriptions of performance thresholds). Additionally, we assessed task engagement time (Figure 7D), quantified as the total time of engagement in the selective detection task (see Methods for assessment details). A 2-way ANOVA test determined significant main effects of region (F(1,44)=11.5, p=<0.01) and hemisphere (F(1,44)=5.21, p=0.03), with target-aligned DLS inactivations displaying the greatest reductions in task engagement.

To better understand the contributions of each site to task performance, we assessed effects of inactivations on hit rate, false alarm rate, behavioral d-prime (separation between hit rate and false alarm rate), and criterion (tendency to respond on target and distractor trials) during the 1-hour selective detection task (Figure 8). For all performance measures, we observed significant main effects of region (p=<0.01), with additional main effects of hemisphere for hit rate and criterion (p=0.03 and p=0.02, respectively). Overall, DLS inactivations resulted in lower hit rates, lower false alarm rates, poorer target-distractor discrimination (d’), and reduced tendency to respond (c). For each measure, the largest impairments were observed for target-aligned DLS inactivations. Moreover, for each measure, target-aligned DLS inactivation sessions were significantly different than non-injected control sessions (task engagement time: control, 5460 +/− 270 seconds; tDLS, 1510 +/− 470 seconds, 72% reduction, p=<0.01; hit rate: control, 0.71 +/− 0.03; tDLS, 0.25 +/− 0.11, 65% reduction, p=<0.01; false alarm rate: control, 0.16 +/− 0.01; tDLS: 0.06 +/− 0.02, 63% reduction, p=0.019; d-prime: control, 1.91 +/− 0.11; tDLS, 0.56 +/− 0.31, 71% reduction, p=<0.01; criterion: control, 0.13 +/− 0.08; tDLS, 1.43 +/− 0.30, 11 fold increase, p=<0.01). These data identify target-aligned DLS, rather than MC, as most critical for responding to task-related stimuli.

Interestingly, we found that muscimol inactivation did not invariably reduce response rates. Inactivations of MC resulted in increased false alarm rates compared to control sessions (Figure 8B), which was statistically significant for distractor-aligned MC inactivations (control, 0.16 +/− 0.03; dMC, 0.29 +/− 0.05, 81% increase p=<0.01). For multiple measures, effects of target-aligned DLS inactivation vs distractor-aligned MC inactivation were significantly different from each other and opposite in direction compared to control sessions. This includes hit rate (tDLS, 0.25 +/− 0.11; dMC: 0.82 +/− 0.07, p=<0.01), false alarm rate (tDLS, 0.06 +/0.02; dMC, 0.29+/− 0.05, p<0.01), and criterion (c) (tDLS, 1.43 +/− 0.30; dMC, −0.33 +/− 0.20, p=<0.01). These findings highlight the divergent functional contributions of these two regions on task performance, with target-aligned DLS dominantly contributing to sensory selection and distractor-aligned MC dominantly contributing to distractor response suppression.

## Discussion

In this study, we compared motor cortex (MC) and dorsolateral striatum (DLS) contributions to performance in a learned selective detection task. First, we identified the S1-project sites within MC and DLS as our target regions (Figure 2). Next, we demonstrated that both regions display robust sensory encoding (Figure 3 and 4) as well as motor-related and choice-related signals (Figures 3, 5, 6) during expert task performance. Lastly, targeted pharmacological inactivation studies identified DLS as essential for responding to task-related stimuli (Figures 7, 8). Collectively, our data support DLS, and not MC, as a crucial bottleneck in linking task-related stimuli to prepotent responses in this learned task (Figure 9).

**Figure 9.**
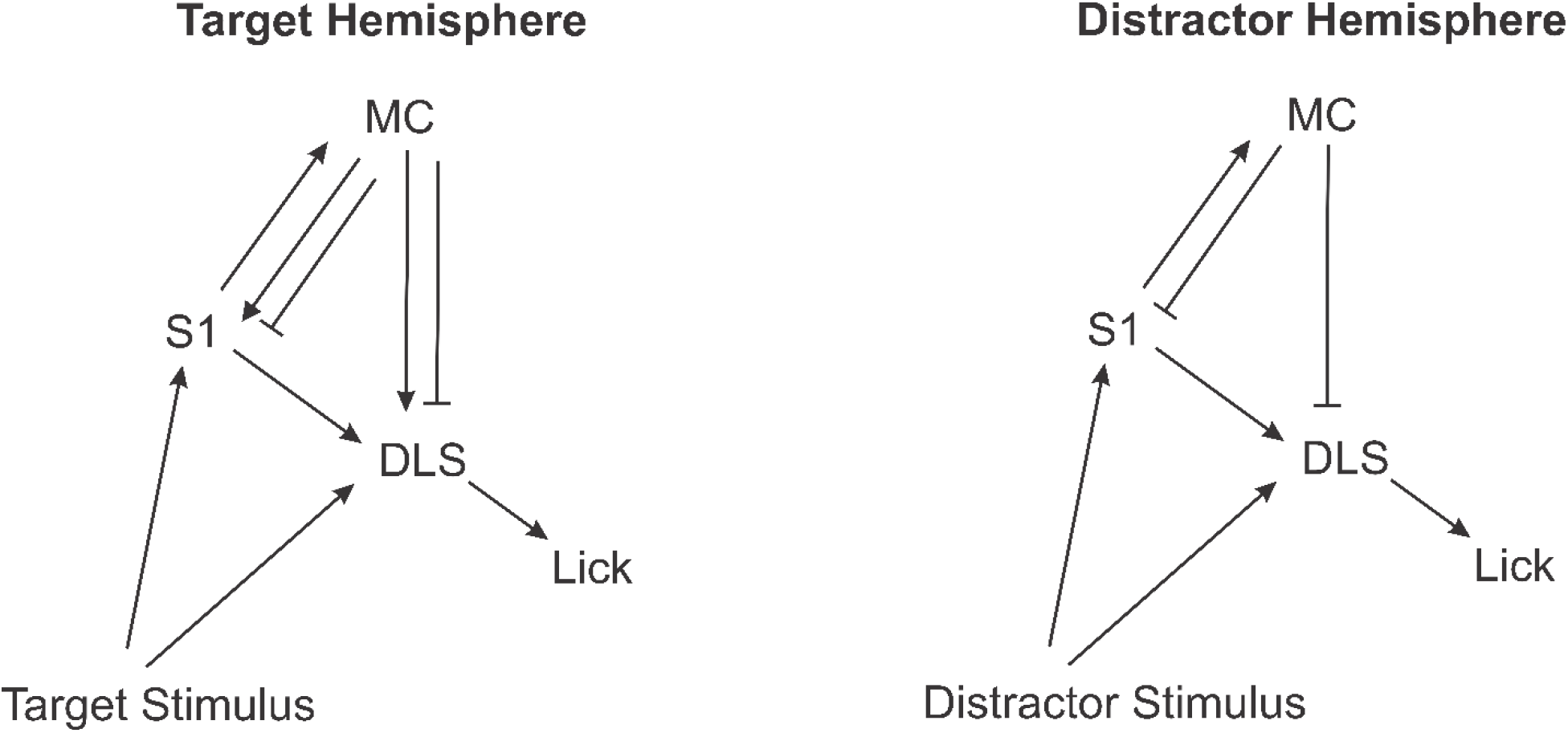
Functional circuit model for how MC and DLS interact to implement sensory selection. Arrows depict net excitatory pathways whereas blunt arrows depict net inhibitory pathways. In this model, target-aligned DLS functions as a critical node in transforming sensory inputs into motor outputs, potentially integrating sensory evidence from S1, MC, and subcortical regions. In contrast, MC acts primarily to modulate the sensory-to-motor transformations that occur through the DLS pathway.

One important consideration when interpreting inactivation studies, particularly for Go/NoGo tasks, is that reductions in task performance may reflect global deficits in motivation, sensation, or response execution rather than specific deficits in task-related processing (Carandini & Churchland, 2013). We interpret the robust impairments during DLS inactivation as reflecting specific deficits in task-related processing for multiple reasons. Most importantly, during the same behavioral sessions, mice with DLS inactivations responded at high rates to auditory-cued reward delivery (Figure 7B), ruling out severe global deficits. Additionally, we observed robust post-stimulus (sensory) and pre-response (possibly motor command) signals within DLS, as would be required for participation in sensory-motor transformations. Whether the actual sensory-to-motor transformation is computed within DLS or is computed elsewhere and propagated to DLS remains unknown. However, our data support DLS as an indispensable node in the sensory-motor transformation process in this task.

Our findings support current frameworks for the importance of DLS in learned tasks (Atallah et al., 2007; Dhawale et al., 2021). An unexpected finding, however, is that target- aligned DLS inactivations did not only reduce responding to target stimuli (Figure 8A), but also robustly reduced responding to distractor stimuli (Figure 8B). We speculate that through learning, response triggering for target and distractor stimuli become conditioned on the activation of target-aligned DLS. Understanding the neural mechanisms underlying this conditioned response triggering, and its possible dependence on dopamine neuromodulation (Gerfen et al., 1990; Gerfen & Surmeier, 2011; Kravitz et al., 2010; Surmeier et al., 1996) are important topics of future research.

It was recently proposed that baseline (prestimulus) activity within the striatum may regulate task engagement, with higher prestimulus activity priming motor execution by bringing the network closer to response threshold (Steinmetz et al., 2019). This framework would have predicted above chance prestimulus choice probability for target-aligned DLS. In contrast, we find that bilateral DLS, like the rest of dorsal neocortex (Marrero et al., 2021), displays below chance prestimulus choice probability in response to target stimuli (Figure 6E). This difference may be due to sensory modality or task design. Nonetheless, our data suggest that for both cortical and subcortical structures within the whisker system, noise reduction enhances stimulus detection.

An important open question is which basal ganglia outputs are most relevant for response triggering. The basal ganglia may project back to regions of motor/frontal cortex not studied here, which may ultimately signal the motor command. And yet, there is growing appreciation of direct projections from the basal ganglia to subcortical motor structures, including the superior colliculus and other brainstem motor structures (Lee & Sabatini, 2021; Saitoh et al., 2003; Takakusaki et al., 2003; Takakusaki et al., 2011). We speculate that, at least in the rodent, striatal selection does not require motor cortex for response execution.

In comparison to DLS, our findings indicate that MC is less critical for responding to task-related stimuli, despite this region demonstrating robust sensory and early choice-related signals. We interpret this discrepancy between representation and function in MC as due to redundant sensory-motor processing through the DLS. Recent studies have shown that motor cortex is required for skill learning, yet is dispensable for performing well-learned tasks (Hwang et al., 2021; Kawai et al., 2015). Thus, it is possible that sensory-motor processing through MC is more important during learning than during expert performance as tested in this study.

Although our data support that MC is not critical for sensory detection in expert mice, we do find evidence for MC involvement in distractor response suppression, especially for distractor-aligned MC (Figure 8B). This finding is consistent with a growing literature, primarily in rodents, suggesting essential roles of motor cortices in suppressing prepotent responses (Ebbesen & Brecht, 2017; Murakami et al., 2017; Zagha et al., 2015). Further studies are required to determine the mechanisms underlying this function.

We do recognize important limitations of our study. Foremost, the multiunit activities analyzed here represent the summed spiking outputs of neuronal ensembles consisting of different cell-types. While we do observe systematic differences in the dynamics of pre-response and choice-related signals in MC and DLS, we recognize that there is likely a much larger heterogeneity of responses of single units. Second, our pharmacological inactivations do not distinguish between the multiple output pathways that may be responsible for the consequent behavioral effects. Follow-up studies with cell-type resolution and pathway-specificity will provide many important mechanistic insights into how DLS and MC contribute to sensory selection.

## Acknowledgements

This work was support by the National Institutes of Health Grant R01NS107599 (to E.Z.). We thank Trevor Zimmerman-Thompson and Emaan Kaur for helping with training of mice. We are also thankful to Zhaoran Zhang, Krista Marrero and Krithiga Aruljothi for their feedback in all stages of the project.

